# Differential interference with actin-binding protein function by acute Cytochalasin B

**DOI:** 10.1101/2024.09.11.611976

**Authors:** Christopher Lambert, Marius Karger, Anika Steffen, Yubo Tang, Hermann Döring, Theresia E. B. Stradal, Pekka Lappalainen, Jan Faix, Peter Bieling, Klemens Rottner

**Author notes:** <u>Author for correspondence:</u>, or.

## Abstract

Dynamic actin filament remodeling is crucial for a plethora of fundamental cell biological processes, ranging from cell division and migration to cell communication, intracellular trafficking or tissue development. Cytochalasin B and -D are fungal secondary metabolites frequently used for interference with such processes. Although generally assumed to block actin filament polymerization at their rapidly growing barbed ends and compete with regulators at these sites, our molecular understanding of their precise effects in dynamic actin structures is scarce. Here we combine live cell imaging and analysis of fluorescent actin-binding protein dynamics with acute treatment of lamellipodia in migrating cells with cytochalasin B. Our results show that in spite of an abrupt halt of lamellipodium protrusion, cytochalasin B affects various actin filament barbed end-binding proteins in a differential fashion. Cytochalasin B enhances instead of diminishes the accumulation of prominent barbed end-binding factors such as Ena/VASP family proteins and heterodimeric capping protein (CP) in the lamellipodium. Similar results were obtained with cytochalasin D. All these effects are highly specific, as cytochalasin-induced VASP accumulation requires the presence of CP, but not *vice versa*, and coincides with abrogation of both actin and VASP turnover. Cytochalasin B can also increase apparent barbed end interactions with the actin-binding β-tentacle of CP and partially mimic its Arp2/3 complex-promoting activity in the lamellipodium. In conclusion, our results reveal a new spectrum of cytochalasin activities on barbed end-binding factors, with important implications for the interpretation of their effects on dynamic actin structures.

## Introduction

The polymerization of actin monomers into filaments is a key reaction for cytoskeleton assembly in eukaryotic cells, and paradigmatic for how dynamic, polymeric assemblies generate forces within and between cells and tissues. ^1^ Dynamically assembled actin filaments and the complex actin structures they form arise from the collective biochemical activities of a plethora of actin-binding proteins, which in turn are subject to regulation by the mechanics and geometry of the actin networks they construct. ^2^ Details on such regulatory feedbacks, however, are just beginning to be elucidated. An efficient method to address the role of actin dynamics in the various distinct structures formed in cells and tissues^3,4^ constitutes cell treatment with plasma membrane-permeable compounds interfering with filament assembly. The most frequently used compounds that interfere with actin polymerization are latrunculins, which sequester the building blocks of polymerization, i.e. actin monomers, and cytochalasans, which target the fast growing, so-called barbed ends of actin filaments to inhibit their elongation. ^5,6^

One of the most prominent and best-studied protrusive structures powered by active actin filament polymerization is the lamellipodium, which is crucial for mesenchymal cell migration on flat, rigid substrates, ^7^ and re-purposed in related processes, such as micropinocytosis, phagocytosis or pathogen uptake. ^8^ In all these structures, the barbed ends of actin filaments are polymerizing while abutting the tip of the protruding membrane, hence exerting pushing forces. These forces translate into a combination of forward membrane protrusion and rearward actin network flow. ^9^ Interestingly, the pushing energy exerted this way can also directly affect the biochemical activities and turnover of the barbed end-binding proteins residing at these sites. ^10^ However, the potential impact of cytochalasans on the regulation and dynamics of such barbed end regulators remains unclear despite its necessity for understanding the phenotypic outcomes of such treatments. Here, we provide a systematic characterization of the effects on distinct actin-binding proteins of acute, lamellipodial treatments with the most popular cytochalasins in the literature, cytochalasins B and -D (CB and CD). Our results, amongst other insights, reveal the counterintuitive, lamellipodial accumulation of various barbed end binders, such as the Ena/VASP family of actin polymerases, upon cytochalasin-induced inhibition of actin polymerization and the surprising dependence of this effect on the Ena/VASP antagonist CP.

## Results and Discussion

### Cytochalasins B and D increase accumulation of VASP at the lamellipodial leading edge

Seminal studies more than two decades ago on Ena/VASP-family actin filament polymerases have established their positive role in actin-based lamellipodial protrusion. A prominent example of this constitutes Ena/VASP inhibition by mitochondrial sequestration (Bear et al., 2002), ^11^ which confirmed the correlation between their accumulation in lamellipodia and the efficiency of forward protrusion. ^12^ Bear and colleagues described VASP reduction upon low dose treatment with cytochalasin D (CD), which spawned the so-called displacement model, which posits that CD outcompetes VASP from lamellipodial barbed ends. In addition, it was suggested that lamellipodial VASP localization is mediated by its association with filament ends. ^11^ However, both of these conclusions remain controversial. ^13,14^ Here we revisited our preliminary observation that blockade of lamellipodial protrusion by high concentrations (52 µM in the application needle) of locally applied cytochalasin B (CB) increases rather than decreases lamellipodial VASP accumulation, ^14^ which occurs acutely and in a fully reversible fashion (Figure 1A-C, Video S1). Interestingly, recovery from the treatment occurs by rearward movement and continuous dissipation of the former lamellipodial front (orange arrowheads in Figure 1A, B at different time points) and its replacement by a new protruding VASP front (light blue arrowheads in Figure 1A, B, Video S1). Moreover, we observed a highly similar response to acute treatment with high dose cytochalasin D (CD, 25 µM), even more dramatic in extent (Figure 1A-D, Video S2), but less reversible than with CB (Figure 1A-C). This is consistent with the previously observed increased potency of CD inhibiting actin assembly compared to CB (up to 10-fold). ^15^ Importantly, low dose CD did indeed cause a decrease in signal intensity of VASP at the lamellipodium tip (Figure 1A-C, and for quantification Figure 1D), as described in fixed cells before. ^11^ This, however, did not coincide with complete abrogation of lamellipodial protrusion, inconsistent with the displacement model, and was accompanied by significant widening of the lamellipodial VASP signal, unlike treatment with high dose CD (Figure 1E). Video microscopy revealed this low dose widening to result from capture and rearward movement of former tip-associated VASP molecules (Video S3), likely driven by incompletely inhibited actin network polymerization at the tip. Together, all these data suggest that the suppression of barbed end elongation of lamellipodial actin filaments by cytochalasins (B or D) coincides with their increased instead of decreased association with Ena/VASP family proteins.

**Figure 1.**
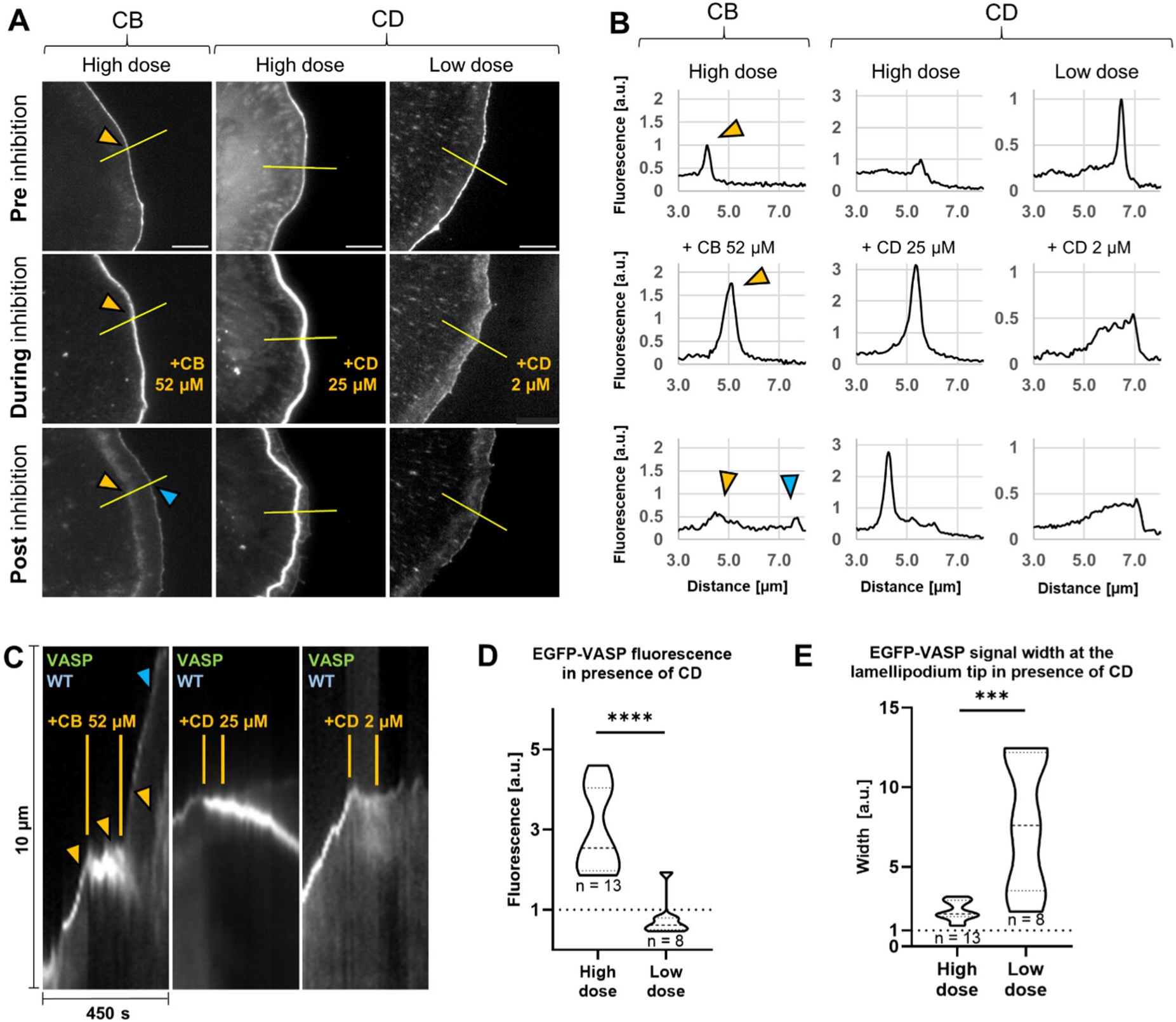
Response of EGFP-VASP dynamics to local applications with CB and CD. (**A**) B16-F1 wild-type (WT) cells transiently expressing EGFP-tagged VASP and imaged before (Pre), during and after (Post) inhibition of protrusion by local application of CB or CD, as indicated. Orange arrowheads in left panels depict VASP signals at lamellipodium tip pre and during inhibition (enhanced in the latter), and the remaining, accumulated signal at the former leading edge post inhibition. The blue arrowhead depicts VASP accumulation at the newly formed lamellipodium tip upon CB washout. Yellow lines show respective positions of the leading edges subjected to line scan and kymograph analyses shown in B and C. Scale bars correspond to 5 µm. (**B**) Line scans of EGFP-VASP signal normalized to the maximum signal preceding local application. Fluorescence plots correspond to lines in A from left to right, and colored arrowheads in left panels to relative signal intensity peaks at respective lamellipodial positions marked in A. (**C**) Kymographs (space-time plots) of EGFP-VASP signal in WT cells along yellow lines drawn in A, depicting 10 µm of line length displayed on the ordinate (kymograph bottom corresponds to line end inside the cell) over a time period of 450 seconds, as indicated. Orange lines mark time periods of active local application of respective cytochalasin to the leading edge. Colored arrowheads again correlate with arrowheads shown in A and B. Color codes for labeling at panel top: green, EGFP-tagged protein and blue, cell type. (**D**) Quantification of maximal EGFP-VASP fluorescence signal and (**E**) signal width following local CD exposure in conditions, as indicated, and normalized in each case to frames preceding treatment start. Data displayed as truncated violin plots (see Methods), n: number of analyzed, individual cells; for statistics see Methods.

### CB treatment differentially affects VASP and FMNL2 and reduces VASP turnover at the lamellipodium tip

Given the surprising effect of cytochalasins on distribution and dynamics of VASP, we wondered whether this behavior would be shared by additional actin filament polymerases, such as formins. Due to the more complex and less reversible VASP responses to CD treatments (Figure 1), subsequent experiments were mostly performed with local application of high dose CB, and results systematically analyzed and quantified. As exemplified in Figure 1C, D, a direct comparison of the behavior of given lamellipodial factor was obtained by employing kymography and quantification of the relative change of fluorescence intensity following CB treatment as compared to immediately before treatment (normalized to 1 in each case, see Methods). We first inspected wild-type cells, and found that unlike the increase of leading edge VASP to about 150%, the increase of the FMNL formin family member FMNL2, which contributes to lamellipodium protrusion downstream of the Rho-GTPase Cdc42 (Block et al., 2012), ^16^ was marginal (Figure 2A, a and Video S4, top panels). As opposed to VASP, the signal of FMNL2 fluorescence did also not widen significantly, all indicative of a differential effect on these two distinct actin polymerases. Interestingly, the VASP response to CB treatment was enhanced in a statistically significant fashion as compared to wild-type in cells genetically disrupted for lamellipodial FMNL family proteins, FMNL2 and −3, ^17^ to app. 200% (Figure 2B, b, Video S4, compare top with bottom left panel), suggesting a higher, relative contribution of VASP to the lamellipodial actin network in these conditions. The marginal effect of CB on FMML2 accumulation, however, was not altered by the absence of Ena/VASP proteins (EVM KO, Figure 2C, c, Video S4, compare top with bottom right panel).

**Figure 2.**
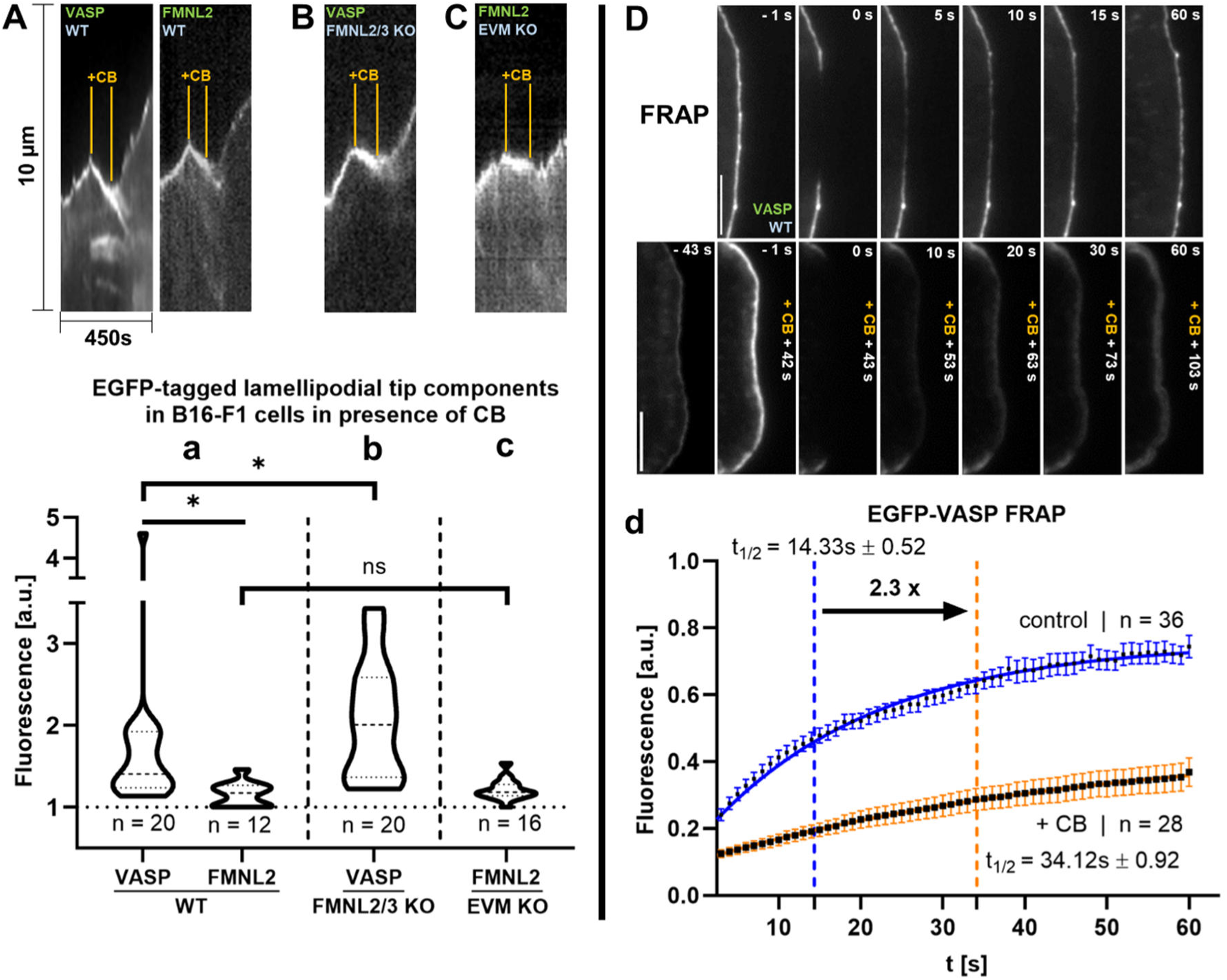
Responses of VASP and FMNL2 to local CB in the context of different genetic backgrounds and CB-induced suppression of VASP turnover at the lamellipodial tip. (**A-C**) Representative kymographs of epifluorescence signals at lamellipodial tips of actively protruding cells transiently expressing EGFP-tagged lamellipodial components (each indicated in green) and in distinct genetic knockouts (cell types indicated in blue), with time windows of CB treatments (local application with high dose, 52µM needle concentration) marked again with orange lines. Note that the onset of CB treatment strictly correlates with abrupt halt of lamellipodial protrusion in each case, frequently followed by modest retraction and robust recovery of protrusion following CB needle removal (end of treatment). For VASP, CB-triggered halt of lamellipodium protrusion is accompanied by the accumulation of fluorescence relative to the intensity before treatment, which occurs less pronounced for FMNL2. (**a-c**) Quantitation of maximal, normalized fluorescence acquired from n individually analyzed cells (for experiments as shown in A-C) and visualized as violin plots (as in Figure 1). (**D**) Lamellipodial fluorescence signals of wild-type cells (WT) transiently expressing EGFP-tagged VASP in the absence (control) or presence of locally applied CB. Labels in horizontal font at panel tops depict time points before and after bleaching, whereas vertical labeling depicts time after CB treatment start, as indicated. Scale bars correspond to 5 µm. (**d**) Corresponding fluorescence recovery curves of EGFP-VASP in the absence (control, blue) and presence (orange) of CB over 60 seconds post-bleaching. Individual data points are arithmetic means and standard errors of means from n analyzed individual cells. Respective half-times of recovery, indicated by colored, dotted vertical lines were calculated upon fitting from solid curves as described in Methods.

The specificity of the VASP response to CB prompted us to explore more directly potential mechanisms behind the observed, reversible VASP accumulation. Localization and dynamics of lamellipodial tip components, like VASP, are controlled by a balance between association to and dissociation from the lamellipodial tip membrane, as visualized e.g. by fluorescence recovery after photobleaching (FRAP). ^18^ Turnover of EGFP-VASP at lamellipodium tips of B16-F1 wild-type cells was previously established to occur with a half time of recovery (t_1/2_) between 11 and 15 seconds, ^18,19^ which well fitted the numbers obtained here in the absence of CB (Figure 2D, d). Interestingly, turnover of some lamellipodial tip regulators such as WRC can be enhanced by active actin polymerization. ^10,20^ Consistently, CB-mediated inhibition of actin polymerization was accompanied by strongly reduced VASP turnover to a measured half time of recovery (t_1/2_) of about 34 seconds (i.e. app. 2.3-fold) as compared to control (Figure 2D, d, see also Video S5). These data suggest that VASP accumulation is the result of suppressed protein dissociation. Structural and biochemical analyses will be required to dissect to what extent reduced actin assembly alone or additional, structural changes at barbed ends are causative of this outcome (see also below). Based on these data, we conclude that VASP dynamics and turnover at the edge of polymerizing, lamellipodial actin networks are highly sensitive to changes in active actin polymerization, likely by increased actin filament association (see above) and thus reduced dissociation into the cytosolic pool.

### VASP molecules display differential responses to CB treatments at lamellipodial *versus* microspike tips

We noticed that while VASP accumulated at the lamellipodium edge following high dose application of CB, it was simultaneously and selectively removed from the tips of microspikes, which then rapidly disassembled (Video S4, top left). This suggests that VASP is selectively trapped and accumulated at the edge of the lamellipodial actin network upon CB, while it is being removed from the tips of embedded microspike bundles. This behavior is reminiscent of previous findings in neuronal growth cones, where selective removal of filopodia was observed following exposure to low doses of CB. ^21,22^ These observations, combined with the notion that genetic removal of all three Ena/VASP family members diminishes microspikes normally embedded into B16-F1 lamellipodia, ^23^ suggest that VASP molecules accumulated at microspike tips rapidly vanish upon CB, which in turn causes their disassembly. Strong sensitivity of microspikes to high dose CB treatment was also apparent from the instantaneous removal of the microspike-enriched bundling protein Fascin (Figure 3A). ^24^ Detailed analysis of Fascin dynamics during CB application and after washout revealed its initial, CB-triggered removal mostly from the rear of these bundles, likely coincident with their disassembly, and re-association upon CB washout from the polymerizing front of the bundle (Figure 3B, Video S6). Interestingly, Myosin X, an unconventional myosin previously implicated in filopodia and microspike formation in various cell types^25,26^ and recently found to operate upstream of Ena/VASP clustering at microspike tips, ^27^ was completely unaffected by local CB application (Figure 3C, Video S6). These data show that the selective VASP removal from microspike tips can be uncoupled from its prominent upstream regulator Myosin X, and more generally, that not all factors involved in the generation of microspikes respond to CB with dissipation. Moreover, even a single actin regulatory factor such as VASP can differentially respond to CB, depending on the context of the actin structure it is active in. This specificity must be due to actin structure-dependent differences, most likely reflecting the presence or absence of other actin assembly regulators and hence differential regulation of barbed end polymerization (see below).

**Figure 3.**
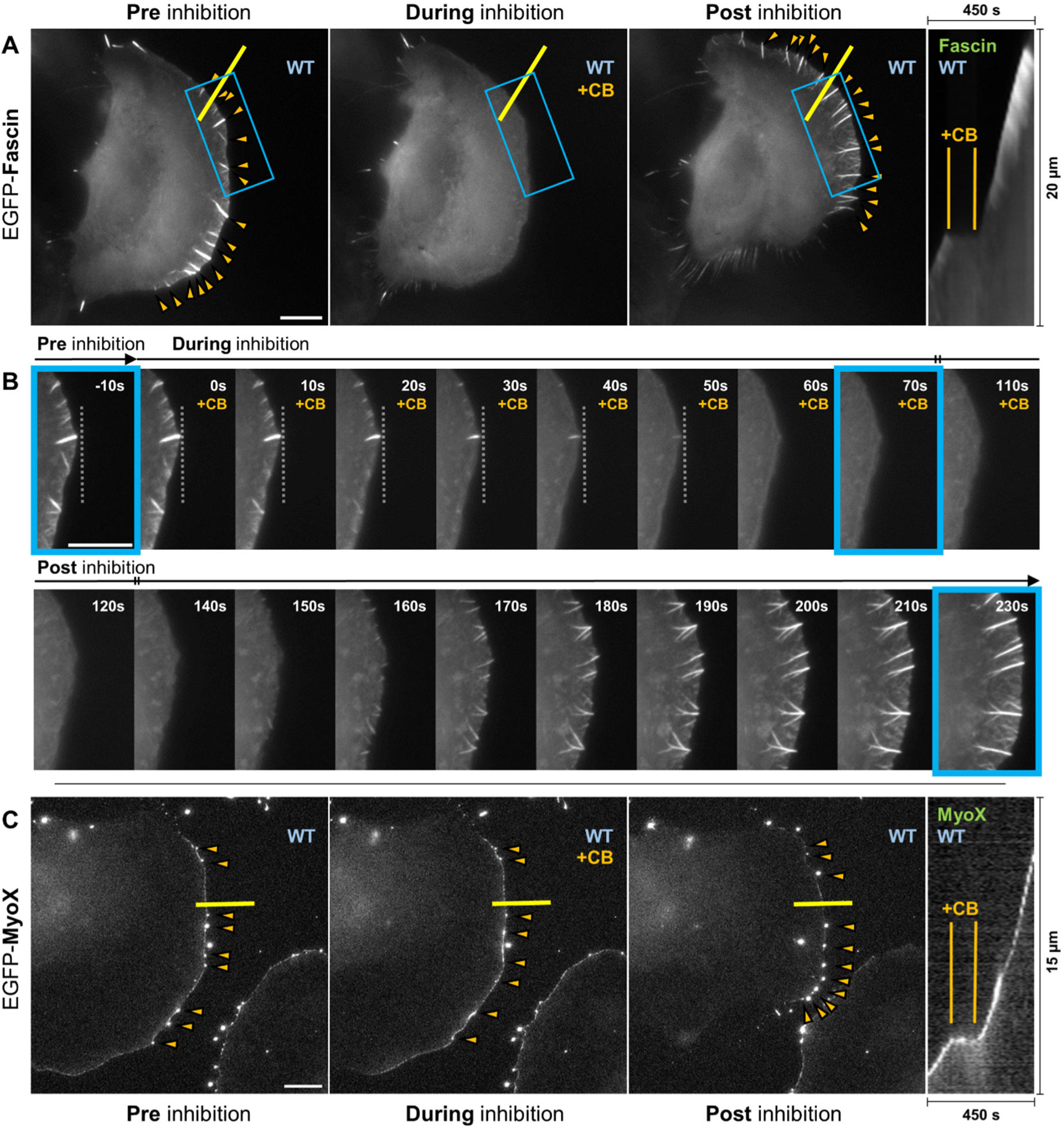
Responses of the actin bundler Fascin and the protrusion regulator Myosin X (MyoX) to local application of CB. (**A-C**) WT cells transiently expressing EGFP-Fascin (**A, B**) and EGFP-MyoX (**C**) before (Pre), during and after (Post) exposition to CB (high dose). Orange arrowheads mark Fascin (**A**) and MyoX (**C**) accumulations throughout the experiment. Note that the Fascin signal shown in (**A**) rapidly vanishes in the presence of CB, while the MyoX signal appears unaffected. Yellow lines indicate positions for kymographs shown to the right in A and C. Kymography as in Figures 1 and 2. (**B**) Individual time lapse panels of insets shown in A, and depicting the signal change during CB treatment (indicated in orange) and recovery at high temporal resolution. Time points displayed in A are additionally highlighted in B by blue rectangles. Note that the EGFP-Fascin signal vanishes from the rear, and not front of microspikes, the relative position of which during treatment start is indicated by the dotted line. Scale bars in A/C and B are 10 µm and 5 µm, respectively.

### CB transforms lamellipodia into less dynamic actin networks harboring short filaments

Due the relevance of VASP on one hand and of active actin polymerization on the other for the efficiency of actin-based protrusion, ^11,12,23^ we wondered how the changes caused by acute CB would affect the lamellipodial actin network and the core machinery for its generation, i.e. the Arp2/3 complex, its activator WRC as well as CP. We first looked at actin itself, and in spite of the induced accumulation of actin polymerases such as VASP (see above), the overall amount of lamellipodial actin filaments present at CB-inhibited leading edges as deduced by the intensity of EGFP-tagged β-actin was largely unchanged (Figure 4A, a). Consistent with the described loss upon CB of Fascin and VASP from microspike shafts and tips, respectively, our video microscopy data also confirmed that these changes coincide with fully reversible disassembly of respective microspike bundles (see Video S7). We also wondered to what extent actin assembly and turnover in these conditions were affected. Of note, although early biochemical studies already observed slow growth of barbed ends even with highest CB and CD concentrations, ^28^ cell biologists commonly assume actin assembly in cytochalasin-treated cells to be completely blocked. This is true at least for conditions in which a given actin-dependent process, such as lamellipodium or growth cone protrusion, is entirely arrested. ^29,30^ However, our results clearly reveal that protrusion arrest is not accompanied by complete elimination of actin filament turnover (Figure 4B, b, Video S7), consistent with the reversibility of CB effects and the potential, obligatory function for cell integrity and viability of low level actin filament remodeling, at least in the cell cortex. ^9^ Notwithstanding this, the suppression of actin filament turnover is still more than 6-fold as compared to control (Figure 4b), implying dramatic deceleration of regulation and dynamics driving lamellipodial actin polymerization.

**Figure 4.**
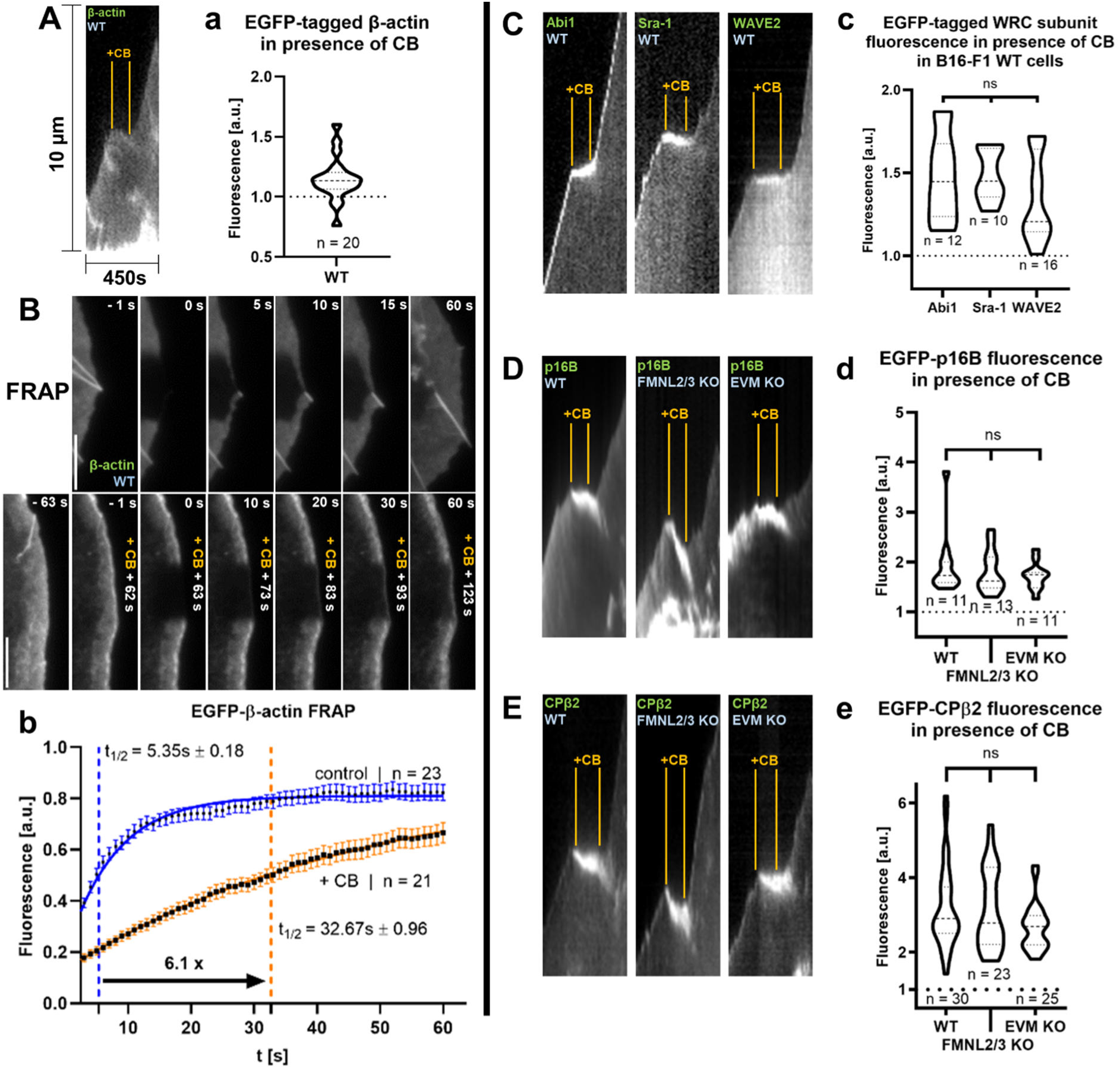
Response of actin dynamics and turnover as well as the lamellipodial actin branching machinery to local CB in the context of different genetic backgrounds. (**A**) Kymography of fluorescently-tagged β-actin in WT cells as indicated and as described in Figure 1. (**a**) Maximal, normalized fluorescence intensities upon CB exposition (high dose) as quantified from n individually analyzed cells in experimental conditions as shown in A and displayed as violin plot. (**B**) Lamellipodial fluorescence signals of WT cells transiently expressing EGFP-tagged β-actin and (**b**) quantitation of its turnover as described for Figure 2D, d. (**C**-**E**) Kymography of cell types and fluorescence of lamellipodial components as indicated and as described for Figures 1 and 2. (**c**-**e**) Quantitation of maximal, normalized fluorescence intensities acquired in each case from n individually analyzed cells (for experiments as shown in C-E), and visualized as violin plots.

Next, we focused on the core machinery of lamellipodial actin nucleation. Protrusion of the lamellipodium requires the continuous activation of the branching activity of Arp2/3 complex, downstream of its main lamellipodial activator, the nucleation-promoting factor (NPF) WRC, followed by activation of the latter through Rac subfamily small GTPases. ^31,32^ Interestingly, although WRC-mediated Arp2/3 complex activation at the leading edge can be circumvented in certain conditions by activation of N-WASP downstream of the Rac-related Rho-GTPase Cdc42, ^33^ WRC recently emerged to be essential for lamellipodial targeting of VASP. ^13^ We tested three subunits of the heteropentameric WRC, all of which target to the tips of protruding lamellipodia, the Abl family kinase interactor Abi1, Sra-1 constituting the interaction surface for Rac GTPases within WRC as well as the ubiquitously expressed WAVE isogene WAVE2. ^34–36^ All of them displayed an increased lamellipodial accumulation upon CB (Figure 4C, c, and as representative example Abi1 in Video S8), similarly in extent to VASP, suggesting a commonality in response to CB between VASP and the WRC/Arp2/3 machinery of branched actin filament assembly. However, this does not yet prove WRC accumulation to be responsible or sufficient for driving CB-induced VASP accumulation (see below).

To explore whether the observed WRC accumulation also translated into increased Arp2/3 complex incorporation at the leading edge, we assessed the changes in fluorescent intensities of the p16B-subunit (ArpC5L) of the complex upon local CB, and found a substantial rise (Fig. 4D, d, Video S9, left panel) to almost 2-fold as compared to before treatment. Considering our results with EGFP-tagged β-actin (Figure 4A, a, Video S7), we conclude that CB-triggered Arp2/3 complex accumulation at lamellipodial edges cannot thus be efficiently translated into actin assembly, likely through rapid interference with filament elongation (Figure 4B). This response was unchanged in cells lacking FMNL family formins (2 and −3), which makes sense considering that these formins promote lamellipodial protrusion largely uncoupled from Arp2/3 complex activity (Figure 4D, d, Video S9, middle panel). ^17^ More surprisingly, identical results were obtained for EVM KO cells lacking all Ena/VASP family members, which are known to display increased average Arp2/3 complex levels in lamellipodia as compared to wild-type cells in the first place, but this did not affect the relative, CB-triggered increase in Arp2/3 complex incorporation (Figure 4D, d, and Video S9, right panel). All these data suggest that the exposure to CB increases the Arp2/3 complex levels in lamellipodia relative to actin, suggesting a relative increase of pointed ends, branches and/or shorter filament stubs, consistent with the inhibition of actin polymerization effected by CB. It can also be deduced that CB, instead of removing lamellipodial actin networks, transforms them into networks with higher densities of shorter filaments labeled at their pointed ends with increased, average amounts of Arp2/3 complex (see model below).

The turnover of lamellipodial actin networks is regulated by a delicate balance between barbed and pointed end binding activities, so we wondered how increased pointed and thus also barbed end numbers would affect the behavior of prominent barbed end binding factors. We already know that Ena/VASP family polymerases were not displaced by CB (see above), but what about CP, the high affinity barbed end interactor crucial for lamellipodia protrusion, the efficiency of Arp2/3-dependent actin assembly and the balance between lamellipodial networks *versus* bundles? ^37–41^ Of note, VASP and CP had already long been attributed an antagonistic relationship at actin filament barbed ends. ^11,42^ Specifically, genetic disruption of all Ena/VASP family members (EVM KO) did not abolish lamellipodia, but decreased their actin network density while increasing the accumulation of CP as compared to wild-type cells. ^23^ Our acute CB treatments performed here revealed interesting analogies to EVM KO cells. Indeed, CB triggered the most dramatic accumulation of lamellipodial CP (almost 3-fold) seen with any actin regulator tested, suggesting a dramatic, relative increase in CP-bound barbed ends, which was independent though (like for Arp2/3 complex) of the presence of both FMNL formins and Ena/VASP family proteins (Figure 4E, e, Video S10). Together, these data establish that local cytochalasin application acutely and reversibly transforms lamellipodial actin networks into less dynamic actin structures harboring shorter filaments with increased amounts of CP and incorporated Arp2/3 complex at their barbed and pointed ends, respectively. This phenotype is similar albeit not identical to cells deprived of Ena/VASP proteins, as CB treatments do not coincide with reduced actin filament densities and correlate with increased accumulation of both Ena/VASP and WRC (see above), unlike lack of Ena/VASP and unchanged WRC in EVM KO. ^23^

### CB-induced VASP accumulation requires the physical presence of CP

As mentioned above, VASP and CP have been ascribed an antagonistic connection, most evident by the increased CP association with lamellipodia lacking Ena/VASP. ^37^ Since CB treatment causes the simultaneous accumulation of both VASP and CP, and as CP accumulation did not require the presence of Ena/VASP (Figure 4E, e, see also Video S10, right panels), we wondered what would happen in the opposite experiment, i.e. in cells lacking the β-subunit of heterodimeric capping protein (CP KO). ^40^ We found, surprisingly, that the observed VASP accumulation did not occur in CP KO cells. Instead, local application of CB onto CP KO cells caused reduction of EGFP-tagged VASP at the lamellipodium tip within less than a minute and its nearly complete removal by 2 minutes (for representative example see Figure 5A, a, B, Video S11). When quantifying the brightest signal after treatment (average of 20 movies), this did not significantly exceed VASP intensities before treatment (Figure 5C, c), as opposed to the approximately 150% in WT cells (Figure 2a), and the latter phenotype could be fully rescued (for mCherry-VASP) by transient re-expression of full length, EGFP-tagged β2-subunit of CP (Figure 5C, c, Video S11).

**Figure 5.**
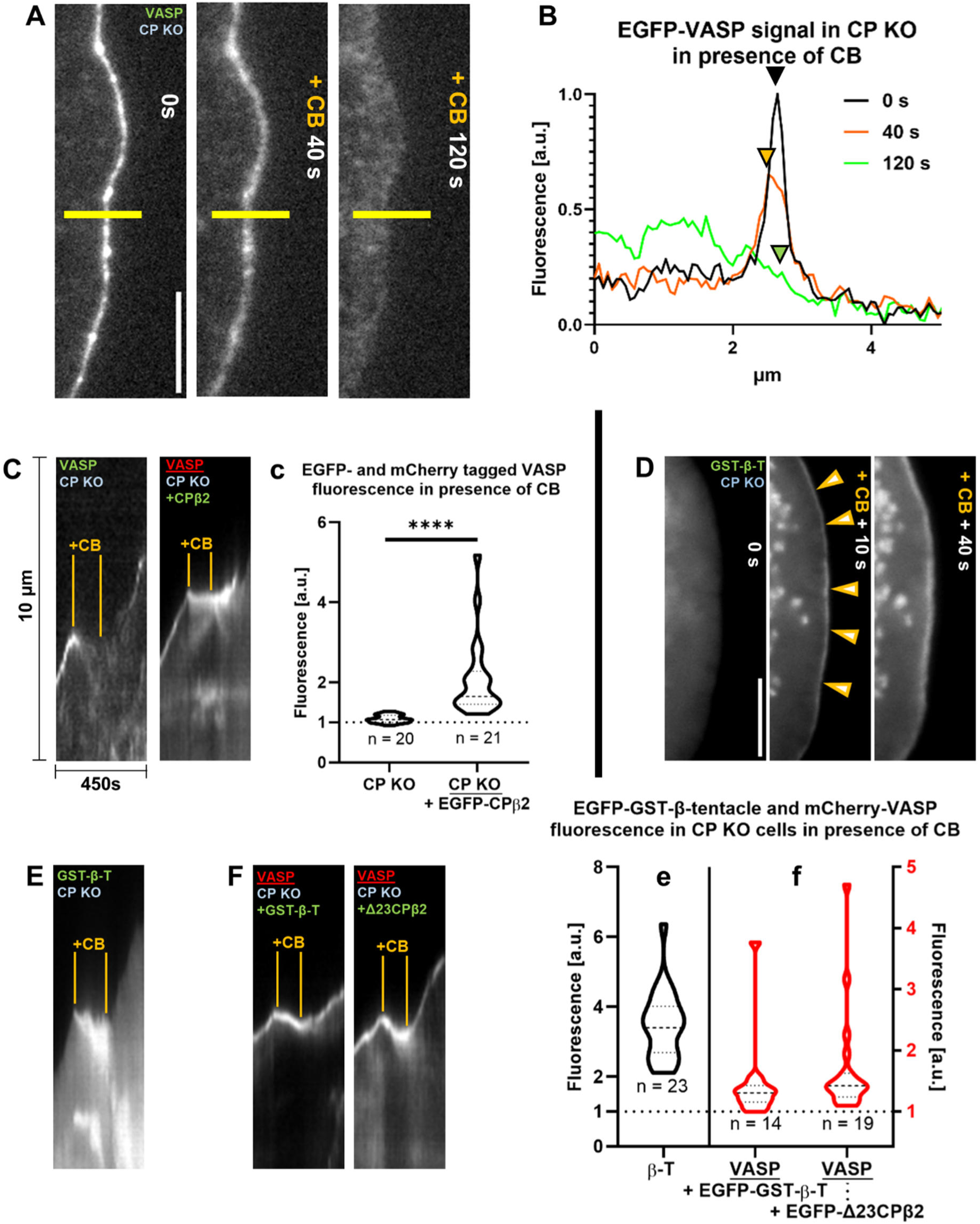
CB-induced VASP accumulation depends on the physical presence of capping protein (CP). (**A**) Signal of EGFP-VASP at the leading edge of CP KO cell exposed to local CB (high dose), as indicated. (**B**) Normalized fluorescence intensities averaged from yellow lines displayed in A. Colored arrowhead tips indicate signals at lamellipodial tips at specific time points upon CB treatment (color-code displayed on the right). Scale bar corresponds to 5 µm. (**C**) Kymographs displayed as described in Figures 1 and 2 and derived from CP null (CP KO) cells expressing fluorescently-tagged VASP with or without reconstituted, full-length CPβ2. Note that EGFP-VASP signal vanishes over the course of CB treatment in the absence of CP (**C**, left), but not if CP function is restored by transient expression of EGFP-tagged, full-length CPβ2 (**C**, right). Color code for expressed protein tags: green, EGFP and red, mCherry (see Methods). (**c**) Maximal normalized fluorescence intensities acquired for n individually analyzed cells with experimental conditions as shown in C. (**D**) Transient expression of an EGFP- and GST-tagged construct of CP’s β-tentacle, described in Methods, in CP KO cells and (**E**) derived kymography. Note that the unspecific lamellipodial signal prior to treatment (0 s) acutely and selectively accumulates at the leading edge (orange arrowhead tips) and other sites of active actin polymerization (ruffles) upon local CB treatment. (**F**) Kymography of mCherry-VASP reconstituted with truncated CP variants as described in C. Labeling color codes as in (C) above. (**e**, **f**) Quantitations of data in experimental conditions shown in E, F for n individually analyzed cells. Violin plots drawn in black and red correspond to the ordinates to the left and right, respectively.

CP is also known to interact with filament barbed ends through several surfaces, but with particular relevance of C-terminal tentacles of both subunits, termed the α- and β-tentacle. ^40,43^ While the β-tentacle is particularly required for efficient, repetitive cycles of NPF-mediated Arp2/3 complex activation, ^40^ it can cap actin filaments even in isolation, albeit weakly, ^43^ and is essential for proper CP accumulation at the lamellipodium tip. ^40^ In spite of this relevant function in lamellipodial CP localization, an EGFP-tagged β-tentacle alone did not target to the lamellipodium tip, not even if dimerized by GST (for construct details see Methods), which had previously enhanced its capping activity *in vitro*. ^43^ Surprisingly, however, if the EGFP-tagged GST-β-tentacle was expressed in the absence of endogenous CP, acute CB treatment led to its transient, but significant accumulation at the lamellipodium tip (Figure 5D, E, e), suggesting that CB binding to barbed ends potentially modified the latter in a fashion that increases capture and binding of the β-tentacle. Interestingly, this effect was not seen in WT cells, likely because endogenous CP effectively competed with the expressed, fluorescent β-tentacle for barbed end-binding (data not shown). This is likely, last not least, due to the 300-fold weaker equilibrium constant for binding the barbed end (*K_cap_*) previously observed for the dimerized β-tentacle. ^43^

Furthermore, we also wondered whether the increased actin barbed end-binding observed by the β-tentacle above would rescue CB-induced VASP enrichment or whether the very same domain would be obligatory for rescue. Interestingly, both the dimerized β-tentacle alone and the β-subunit lacking the β-tentacle (EGFP-Δ23CPβ2) were capable of rescuing CB-induced VASP accumulation (Figure 5F, f).

Thus, the function of CP for promoting Arp2/3 complex activity recently ascribed to the β-tentacle^40^ could be clearly uncoupled from its role in CB-induced, lamellipodial VASP accumulation. In addition, these data suggest that instead of a specific, direct interaction surface on CP for VASP that would keep the latter from rapid dispersal upon CB treatment, barbed end binding by CP or CP fragments (such as the β-tentacle as minimal domain) indirectly impacts on VASP regulation at lamellipodial leading edges. Without this CP function, VASP will be rapidly lost from leading edges treated with CB.

Notably, CB applications in wild-type cells also reduced VASP signals instead of enhancing them in focal adhesions, the latter of which lack CP (data not shown). It is tempting to speculate therefore that differential VASP responses to CB in distinct actin structures are caused by differential amounts of CP present in them. Finally, a similar, accelerated loss of VASP signal from the lamellipodial tip of CP KO cells was also observed upon treatment with high dose CD (Video S12), clearly contrasting the response in wild-type cells (Video S2) and revealing the principal commonality of effects observed for both cytochalasins B and -D (also see Figure 1). All together, we conclude that the cytochalasin-induced enrichments of VASP at lamellipodial leading edges cannot occur in the absence of CP.

### The CP regulator Twinfilin contributes to, but is not directly targeted by CB to promote capping protein accumulation

Recent studies have established that the actin regulator Twinfilin promotes the turnover of lamellipodial actin networks by removing tightly associated CP from barbed ends followed by their depolymerization. ^44^ Since CB treatment caused the accumulation of VASP through CP, we wondered whether this effect required the physical presence of the CP regulator Twinfilin or was modified somehow in cells devoid of this important CP regulator. In cells CRISPR/Cas9-targeted for Twinfilin 1 and −2 (Twf1/2 KO), ^44^ CP that quite significantly enriches close to the edge of the lamellipodium in WT cells, ^18,45^ cannot dissociate from barbed ends and thus localizes throughout the entire lamellipodium, which in turn reduces actin turnover in the latter. ^44^ Interestingly, CB treatment still induced CP accumulation in Twinfilin KO cells, but the extent of this effect was significantly reduced as compared to WT controls (Figure S1A, a). Nevertheless, VASP accumulation upon CB exposure was even more prominent in Twf1/2 KO as compared to WT cells (Figure S1B, b), consistent again with the view that in certain conditions, CP and Ena/VASP family protein accumulation in the lamellipodium can be antagonistic to each other. ^23^ Finally, Twinfilin itself, which weakly associates with the entire actin network throughout the lamellipodium, ^44^ was also enriched by CB in WT cells (Figure S1C, c), probably indirectly through its desperate association with high amounts of CP and/or barbed ends. ^38^ All these data suggest that whereas Twinfilin per definition cannot serve as direct target factor in our experiments essential for CB-induced CP accumulation, it does - as CP regulator - indirectly contribute to CB-induced effects, because its absence causes at least quantitative, statistically significant differences in CP and VASP responses to CB treatments (Figure S1A, a, B, b).

### CB can partially mimic capping protein function concerning Arp2/3 complex activation and actin accumulation

CP was recently shown to promote Arp2/3 complex-dependent actin assembly through interference with non-productive barbed end binding of NPFs such as WRC. Following this reasoning, CP KO cells recently developed in our labs formed lamellipodia with compromised and chaotic protrusion^40^, but those lamellipodia still formed also displayed reduced Arp2/3 complex accumulation (unpublished data). Interestingly, the measured, decreased Arp2/3 complex levels coincided with increased F-actin intensities in these lamellipodia (unpublished data), similar to observations previously published at the total cell level for independently generated CP null cell lines. ^46^ If expressing the Arp2/3 complex subunit p16B (ArpC5L) in CP KO cells, video microscopy revealed weak lamellipodial association of the complex in between multiple, non-Arp2/3-associated microspike bundles (Figure 6A), because the abrogation of CP function is well established to induce bundling and filopodia formation. ^41,46^ Notwithstanding this, acute CB treatment now rapidly transformed this pattern into instantaneous, enhanced Arp2/3 accumulation (Figure 6A), quantified to be significantly higher in CP KO cells than the Arp2/3 accumulation in wild-type cells (Figure 6B, b, left panel). Importantly, this effect was independent of WRC accumulation, as exemplified in this case again by the WRC-subunit Abi1 (Figure 6B, b, middle panels, Video S13, left), since the latter was not additionally enriched in CP KO cells upon CB as compared *versus* wild-type cells. It is tempting to speculate therefore that the CB-induced increase in Arp2/3 complex accumulation seen here is caused by a partial mimicry of a mechanism regularly executed by CP. In spite of CB known to reduce actin filament elongation and the already enhanced F-actin content of CP-less lamellipodia (not shown), a similar enhancement of expressed β-actin accumulation was found in CP KO cells as compared to wild-type, strongly suggesting that the incorporation of Arp2/3 complex was driven by active, polymerization-dependent branching (Figure 6B, b, right panels, and Video S13, right panel). All these data confirm the complexity of changes accompanying CB-induced actin network transformation, with CP constituting a major regulatory player being central to both VASP accumulation and the extent of Arp2/3 complex-dependent actin branching.

**Figure 6.**
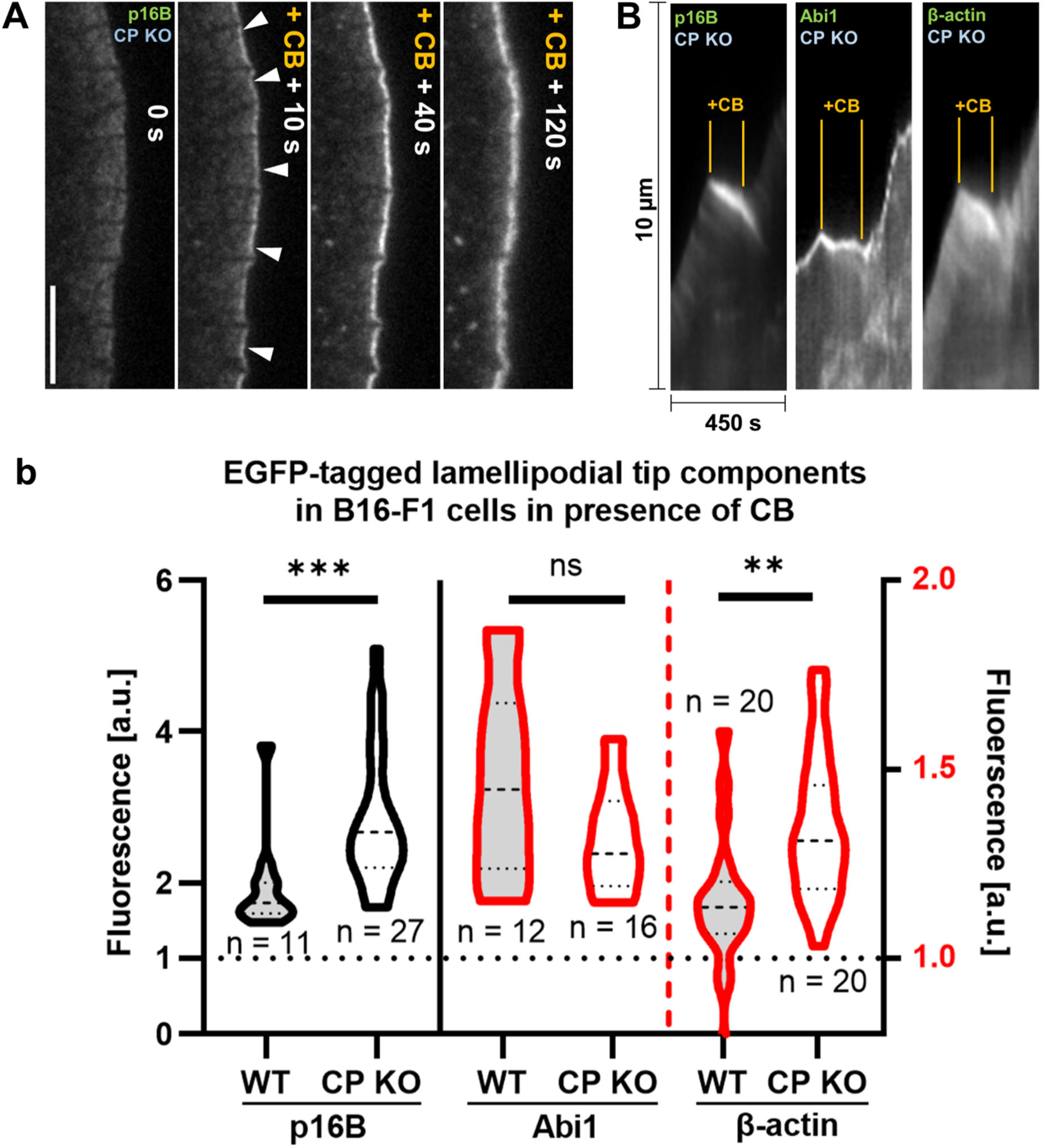
CB-induced enhanced accumulation of Arp2/3 and actin, but not of WRC in the absence of CP compared to WT. (**A**) Transient expression of EGFP-p16B in CP KO cell before and after local application of CB (high dose), as indicated. Scale bar corresponds to 5 µm. (**B**) Kymographs displayed as described for Figures 1 and 2 and derived from CP KO cells expressing fluorescently-tagged p16B (Arp2/3 complex subunit ArpC5L), Abi1 and β-actin. (**b**) Maximal, normalized fluorescence intensities quantified from experimental conditions shown in B for n individually analyzed cells and compared to data obtained with wild-type cells (WT). Grey violin plots display WT data from Figures 4a and 4c, d (left panels) for comparison. Violin plots drawn in black and red correspond to the ordinates to the left and right, respectively.

## Conclusions

We provide evidence for direct effects of cytochalasin (B and D) treatments on actin-binding proteins that are differential in many respects. Firstly, those effects, which are largely reflected by accumulation of actin-binding proteins at sites of active actin polymerization are evident in particular for parts of the Arp2/3 complex-dependent actin assembly machinery. Specifically, this includes the major lamellipodial activator of the latter, WRC, CP, Arp2/3 complex itself as well as Ena/VASP family actin polymerases, but much less so lamellipodial formins such as FMNL2/3. Accumulation of Ena/VASP coincides with suppression (but not elimination) of their turnover as well as of actin assembly, and requires the physical presence of CP. Secondly, described effects are also structure-specific, as Ena/VASP proteins at microspike tips or in adhesions are less prominently affected than those residing at the lamellipodium edge where they co-localize with CP. Thirdly, cytochalasins cause dramatic transformations of highly dynamic actin networks, such as in lamellipodia. The major changes can be deduced as significant shortening of lamellipodial filaments accompanied by increased incorporation of CP/VASP and Arp2/3 complex at barbed and pointed ends, respectively, in wild-type cells (see scheme in Figure 7). These changes are even enhanced concerning actin and Arp2/3 complex incorporation if CB is applied in the absence of CP, whereas VASP accumulation is abolished in these conditions (Figure 7). Our data also allow suggesting a mode of action occurring at the barbed end of lamellipodial actin filaments in wild-type cells upon cytochalasin binding. We propose that CB (or CD) disrupts the antagonistic exchange mechanism of CP and VASP, so that the latter (and perhaps additional factors) are irregularly clamped together at the barbed end in the presence of CB (Figure 7, inset). Consideration of these results will be instrumental for understanding the intricacies of cytochalasin treatments in more complex contexts, such as at the organismal level or if employed as agents against infections and other diseases.

**Figure 7.**
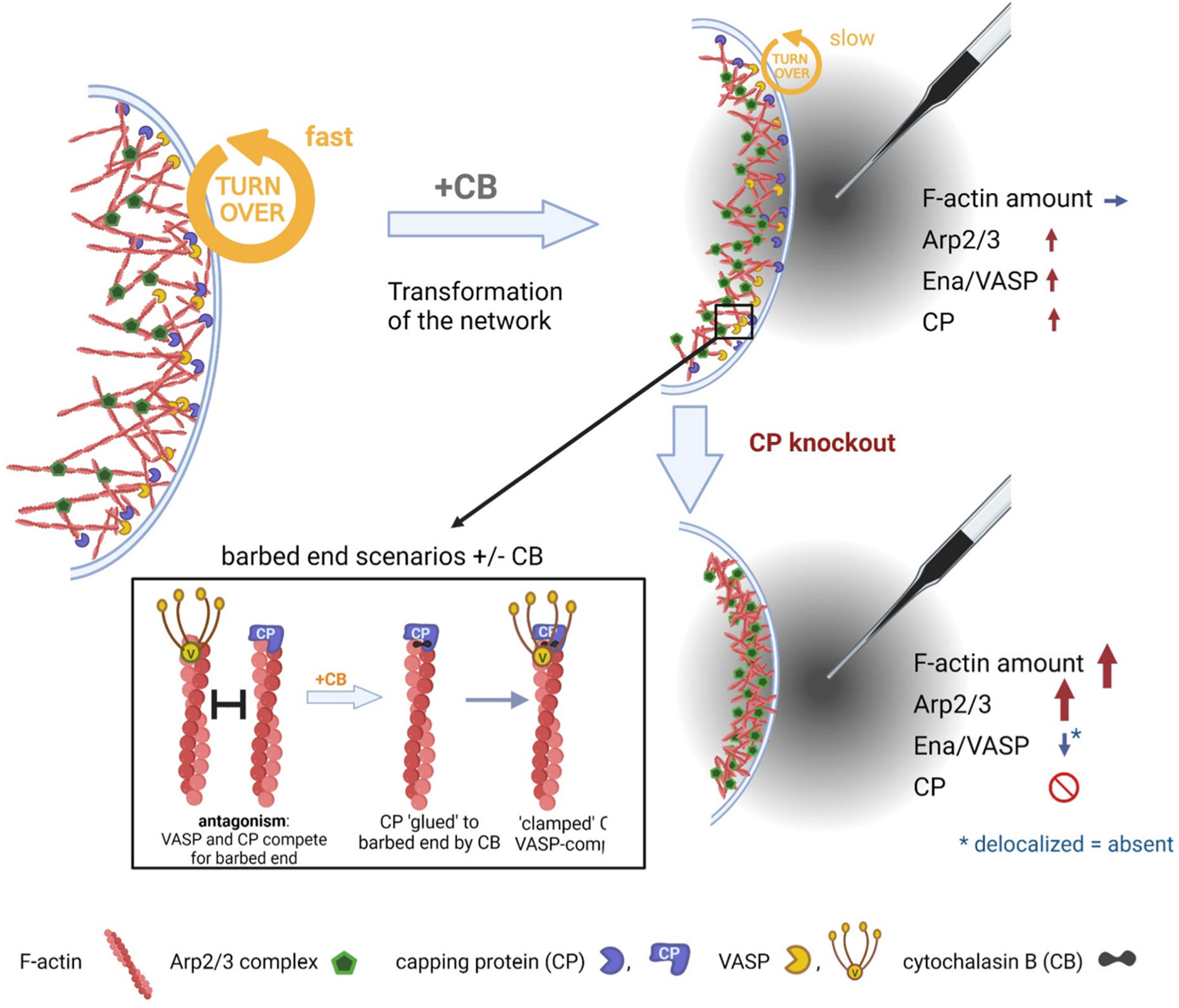
Transformation of lamellipodial actin networks by local CB in the presence or absence of CP. Summary of key observations described in this work upon local, transient treatment of protruding lamellipodia with high dose cytochalasin B (CB, orange flowing out of capillary). Actin filaments and actin regulators are illustrated as described in the legend at the bottom. Note that CB enhances Arp2/3 as well as Ena/VASP and CP accumulations at lamellipodial leading edges while leaving actin levels virtually unchanged. In CP knockout, actin and Arp2/3 intensities are additionally enhanced and Ena/VASP accumulation eliminated. Black rectangle marks a zoom into proposed barbed end populations present at the leading edge of CB-treated wild-type cells. CB transforms mixed populations of VASP- and CP-capped, individual barbed ends present in untreated WT cells (left) into ends glued to CP alone or both CP and VASP clamped together. Structural details are to be elucidated in future efforts.

## STAR*Methods

### KEY RESOURCES TABLE

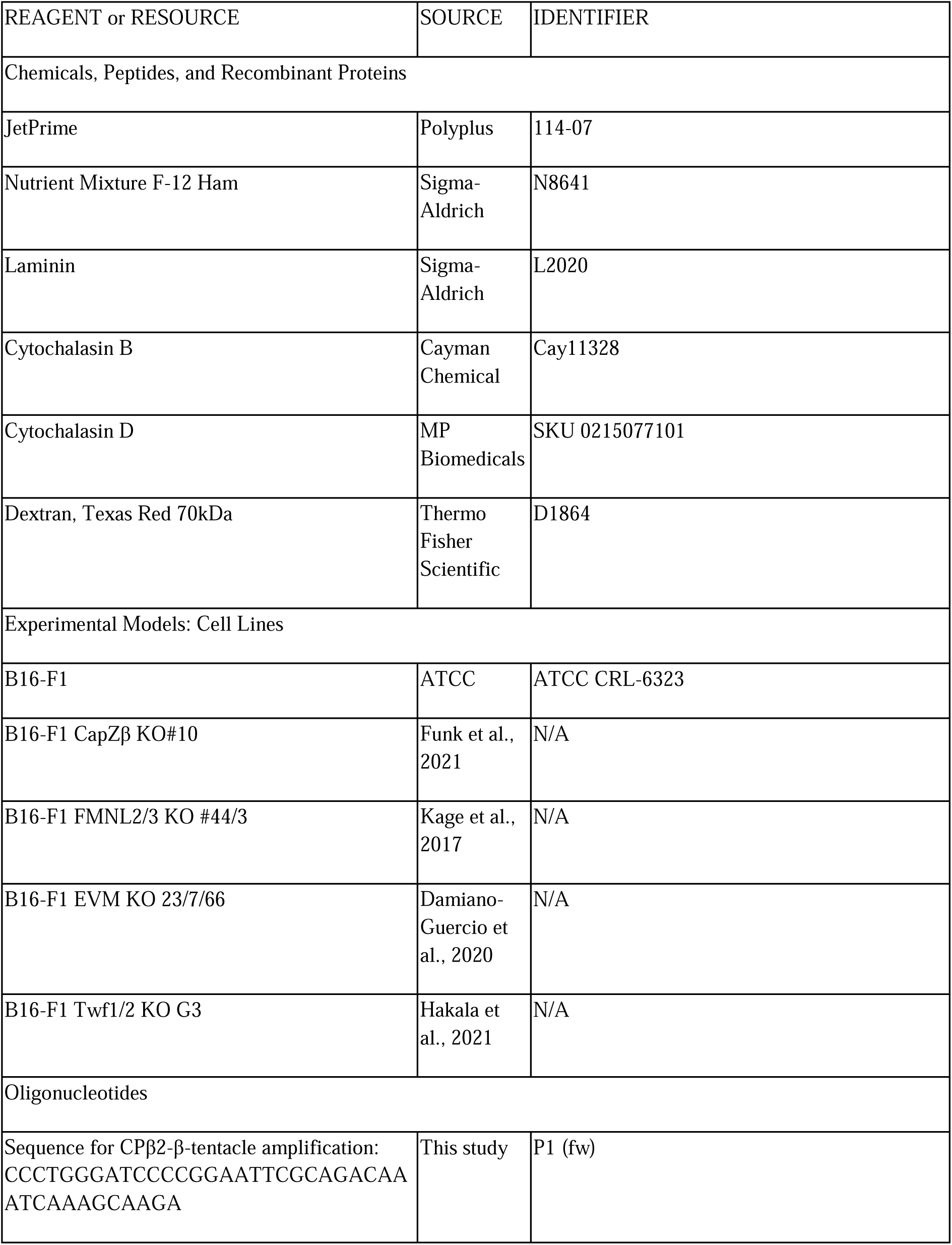
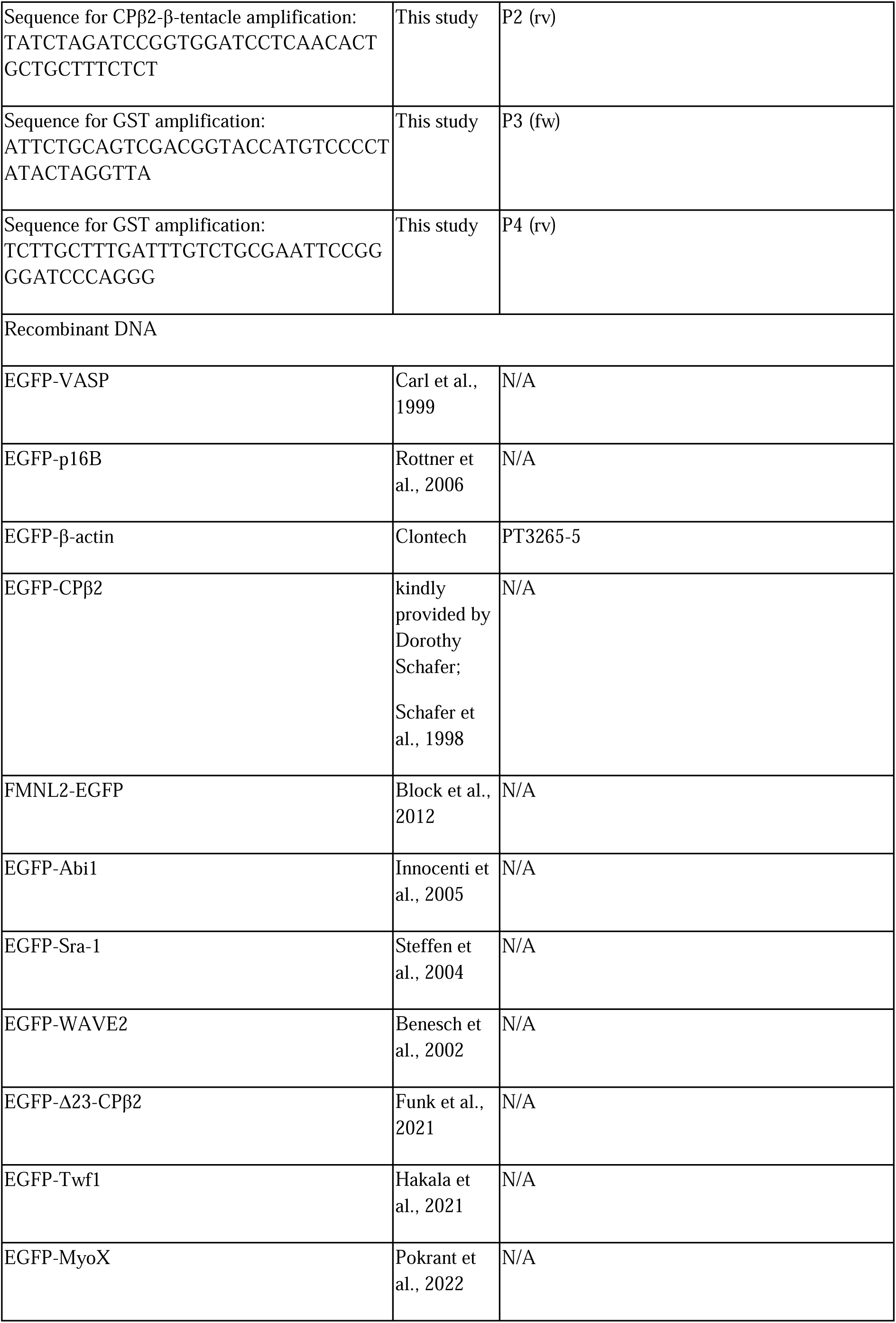
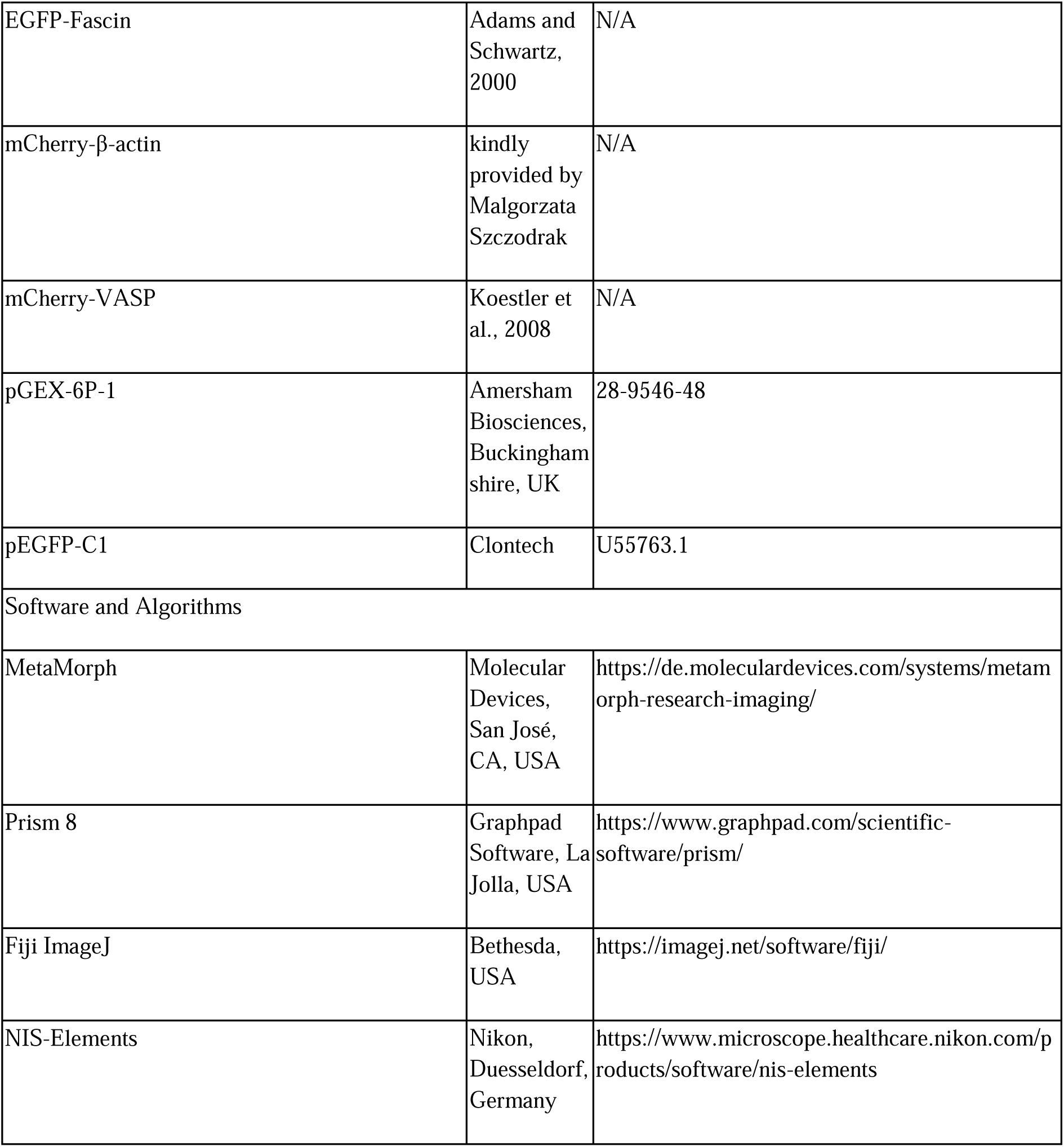

#### Contact for Reagent and Resource Sharing

Further information and requests for resources and reagents should be directed to and will be fulfilled by the Lead Contact, Klemens Rottner.

#### Experimental Model and Subject Details

B16-F1 cells were originally purchased from ATCC (CRL-6323, sex: male) and authenticated by local authorities to be of murine origin and to lack unexpected sequences such as commonly used antibiotic resistance genes or viral sequences. The parental and all genetically-modified, derivative B16-F1 lines are routinely subjected to mycoplasma tests on a biannual basis.

#### Cell culture

B16-F1 mouse melanoma cell line were cultivated in full medium (Dulbecco’s Modified Eagle Medium (DMEM, Life Technologies, Carlsbad, CA, USA) containing 4.5 g/L glucose and supplemented with 200 mM L-glutamine (Life Technologies, Carlsbad, CA, USA) and 10% low endotoxin FCS (Capricorn Scientific, Ebsdorfergrund, Germany) in a humidified incubator at 37 °C, 7.5% CO_2_. Alternatively, a pH-indicator-free and HEPES-buffered microscopy medium containing 10% FCS, 1% L-glutamine and 1% penicillin-streptomycin (10,000 U/mL, Life Technologies, Carlsbad, CA, USA) supplied in F12-HAM (HEPES-buffered medium, Sigma-Aldrich, Missouri, USA) was used as will be indicated further below.

#### Transfections

1 x 10^5^ cells were seeded into 3 cm diameter petri dishes, and next day transfected at 50-70% confluence in single or co-transfection experiments with different plasmids by using 0.5 µg plasmid DNA in total essentially following manufacturer’s recommendations (JetPrime Polyplus Transfection, Illkirch, France; see also ^47^). A total volume of 200 µL was then spread dropwise over the cell culture dish while continuously circling the petri dish. The used plasmids encoding for fluorescently coupled proteins were as follows: EGFP-VASP, ^48^ EGFP-p16B, ^49^ EGFP-β-actin (EGFP-tagged, human β-actin was purchased from Clontech (Mountain View, CA, USA), EGFP-CPβ2 (kindly provided by Dorothy Schafer, ^50^ FMNL2-EGFP, ^16^ EGFP-Abi1, ^51^ EGFP-Sra-1, ^35^ EGFP-WAVE2, ^52^ EGFP-Δ23-CP2, ^40^ EGFP-Twf1, ^44^ EGFP-MyoX, ^27^ EGFP-Fascin, ^53^ mCherry-β-actin and mCherry-VASP. ^54^

#### Construct generation

The EGFP-GST-β-tentacle construct was made as follows: Glutathione-S-transferase (GST) was amplified with pGEX-6P-1 vector (Amersham Biosciences, Buckinghamshire, UK) as template using primers P1 and P2. The β-tentacle of capping protein (nucleotides 736 to 819 of transcript variant 2 of GenBank accession NM_009798.4) was amplified with primers P3 and P4 and the EGFP-CPβ2 plasmid (see above) as template. For primer sequences, see Key Resources Table. The construct was assembled using the SLIC (Sequence- and Ligation-independent Cloning)-method. ^55^ In brief, primers harbored overhangs to the respective neighboring fragment, and resulting PCR products as well as the vector (digested with Kpn1/BamH1) were treated with T4 polymerase/exonuclease and transformed into TOP10 *E. coli* for assembly. The resulting construct was sequence-verified.

#### Live cell imaging

Data were collected using Zeiss or Nikon inverted fluorescence microscopes as follows. For Zeiss, images were recorded in 10 s time intervals by a CoolSnap HQ2 (Photometrics, Tucson, AZ, USA) attached to a Axiovert 135 microscope body, equipped with HXP-120 (Zeiss, Oberkochen, Germany) and halogen lamps for fluorescence and phase contrast microscopy, respectively. Light paths were controlled with Uniblitz shutters (Model D122, Vincent Associates, Rochester, NY, USA) with exposure times ranging from 1-3 s for widefield fluorescence and 100 ms for phase contrast images, using 100×1.3 NA (PLAN-Neofluar, Zeiss, Oberkochen, Germany) or 100x 1.4 NA (PLAN-Apochromat, Zeiss, Oberkochen, Germany) objectives, and EGFP fluorescence excitation and emission visualized using a FITC filter (49002, Chroma, Bellows Falls, VT, USA). Image acquisition was controlled with Metamorph software (Molecular Devices, San José, CA, USA). Heated Warner chambers were mounted and controlled without additional microscope stage heating and humidification.

The Nikon Ti2-E was equipped with a heated, humidified and CO_2_-aerated stage-top incubator with a custom-made lid from Okolab (H301, Ottaviano, Italy), allowing the insertion of a microcapillary into the incubation chamber. Samples were illuminated with a pE-4000 Cool LED light source and excitation and emission visualized with GFP HQ and TRITC filters. Images were acquired using an CFI P-Apo 100x Lambda oil Ph3 1.45 NA objective (Nikon) with exposure times ranging from 100 ms to 400 ms and a pco.edge 4.2 sCMOS camera (Excelitas Technologies, Alzenau, Germany) controlled with NIS-Elements (Nikon, Duesseldorf, Germany). Cells were monitored for the formation of continually protruding lamellipodia, which were exposed to cytochalasins locally applied by gentle but expeditious lowering of a microcapillary for time periods ranging between 100 and 150 s (see below).

#### Local application of cytochalasins

Cells for live cell imaging were transfected and seeded onto coverslips to be mounted on a Warner chamber or into 3.5 cm glass bottom dishes (µ-dish 35 mm, high Glass Bottom, Ibidi, Gräfelfing, Germany) essentially as described. ^47^ Briefly, acid washed glass coverslips and Ibidi dishes were coated with 25 µg/mL laminin (Sigma-Aldrich, Missouri, USA) diluted in laminin coating buffer for 1 h and washed once with 2 mL PBS. Immediately following aspiration, transfected cells were seeded onto coated coverslips in full medium. After allowing them to spread for 3 h, coverslips were mounted into Warner chambers, and the medium replaced with microscopy medium. The latter was exchanged in 1 h time intervals on the microscope to prevent evaporation artefacts. For local application, glass capillaries (Femtotips, Eppendorf, Hamburg, Germany) were filled with cytochalasins pre-mixed with Texas-Red coupled, high molecular weight dextran (70 kDa) in a final concentration of 0.625 mg/mL as inert needle flow and microinjection tracer in microinjection buffer, as detailed earlier. ^56^ Fluorescent dextrane and cytochalasin stock solution were prepared in phosphate-buffered saline (PBS) and DMSO, respectively. Employed, final mixtures contained 2 to 5 µM (for low dose) or 25 µM (high dose) cytochalasin D (MP Biomedicals, Irvine, CA, USA) or 52 µM cytochalasin B (Cayman Chemical, Ann Arbor, MI, USA) in 2.5% DMSO at max. After filling and removal of air bubbles, the capillary was coupled to a FemtoJet or FemtoJet 4i (both Eppendorf, Hamburg, Germany) connected to Zeiss or Nikon inverted microscopes, respectively. After filling and mounting, capillaries were rapidly positioned into the growth medium well above cells routinely applying a constant flow pressure of 30-70 hpa, and with its position controlled by an Injectman NI 2 (Eppendorf, Hamburg, Germany) for the Zeiss or an Injectman 4 (Eppendorf, Hamburg, Germany) for the Nikon microscope. Fluorescent dextrane was omitted in case of co-transfection experiments.

#### Data processing, statistics and illustrations

Fold changes of fluorescent proteins as indicated in the main text were obtained from calculating averaged, maximal fluorescence signal intensities of appropriate regions near lowered microcapillaries upon treatment and divided by averaged signals of respective regions and frames immediately before treatment. Derived, quantified fluorescence intensity data as well as the single width measurements shown in Figure 1E were displayed as so called truncated violin plots, with the width representing data frequency, the three dashed lines separating the quartiles and the middle line representing the median.

All analyses were done using Fiji ImageJ (https://imagej.net, Bethesda, USA). Statistical significance was tested by employing nonparametric Kruskal-Walis and Mann-Whitney Rank sum tests (GraphPad Prism 8.4.3, Boston, MA, USA). Statistical significance was defined with p-values as follows: p ≤ 0.05 = *; p ≤ 0.01 = **; p ≤ 0.001 = ***; p ≤ 0.0001 = ****; n.s., not significant. The scheme in Figure 7 was drawn using BioRender (www.biorender.com).

#### Fluorescence recovery after photobleaching experiments

FRAP movies were recorded using the Nikon Ti-2E equipped with a LUN-F 488/561/640-405 laserbox with separate coupling of a 405 nm laser (50 mW; LUN-F, Nikon, Duesseldorf, Germany) controlled by a FRAP unit (Nikon). Laser intensity was set to lowest possible power to bleach the signal without compromising cell viability. Cells were monitored for continuous lamellipodial protrusion with or without local application of cytochalasin B followed by bleaching of a rectangular field positioned around the lamellipodium of interest. Bleaching following CB treatments was routinely done after 50-70 s treatment periods. Fluorescence recovery was recorded in 1 s intervals for 60 seconds. Total CB treatment durations were limited to a maximum of approximately 130 seconds. Curve fits were calculated in Prism 8.4.3 and half times of recovery calculated from the means of recorded data as described. ^56^ Statistical significance was tested with nonparametric Mann-Whitney Rank sum test.

## Acknowledgments

This work was supported in part by the German Research Council (DFG) through the CytoLabs consortium (DFG Research Unit 5170 individual grants to T.E.B.S. and K.R.) and individual grants Fa330/12-3 (to J.F.) and BI 1998/2-1 (to P.B.) as well as intramural core grants from the Helmholtz Society (to T.E.B.S. and K.R.). C.L. and Y.T. are thankful for stipends granted by the Life-Science Foundation (LSS, Munich) and China Scholarship Council (CSC), respectively.

## Author contributions

C.L. and M.K. performed wet experiments. A.S., Y.T. and H.D. helped with experiments and/or provided essential advice for experiments or data analysis. T.E.S., P.L., J.F. and P.B. provided essential reagents and/or helped with supervision. All authors analyzed and discussed data. C.L., M.K. and K.R. interpreted the data and wrote the manuscript. All authors read and commented on the manuscript and/or amended parts of it.

## Supplementary information

**Figure S1.**
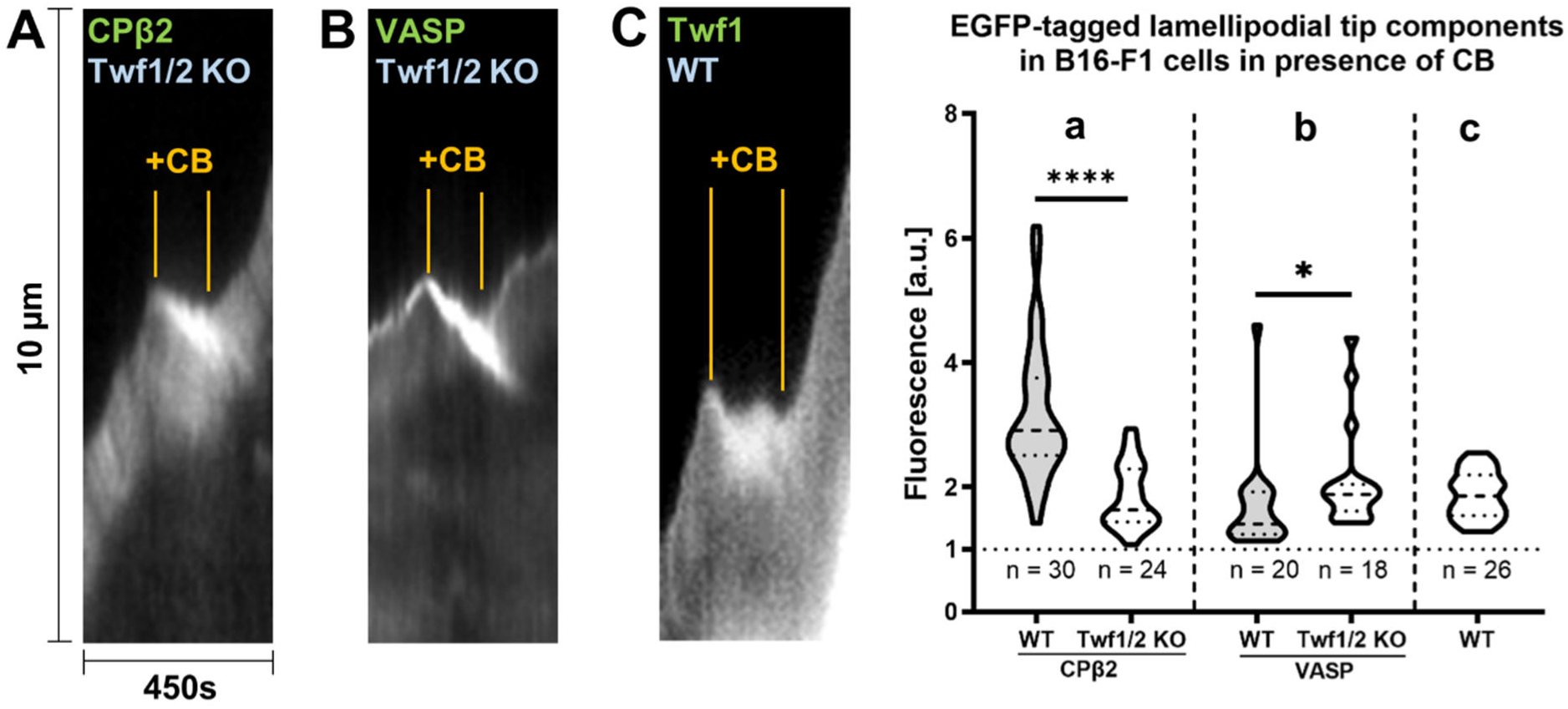
CB-dependent, differential accumulation of CP and VASP in Twf1/2 KO cells. (**A, B**) Kymography of lamellipodial signals of fluorescently-tagged proteins and cell lines as indicated and described for Figures 1 and 2. (**a, b**) Maximal, normalized fluorescence intensities upon CB exposition (high dose) quantified from n individually analyzed cells in experimental conditions as shown in A, B, and displayed as violin plots. Grey violin plots correspond to the data shown in Figure 2a (left panel) and Figure 4e (left panel), and are displayed for comparison. (**C, c**) Representative kymograph (**C**) and quantitation (**c**) of CB-induced accumulation of Twinfilin 1 (Twf1) in WT cells.

**Video S1.** Movie showing representative example of phase contrast and epifluorescence live cell imaging of migrating B16-F1 WT cell expressing EGFP-tagged VASP and subjected to local application of CB (high dose, 52µM needle concentration), as indicated. Local application to the protruding lamellipodium is initiated by lowering of the glass capillary as shown in the phase contrast overview panel on the left, and the duration of active treatment is also indicated (+CB) in yellow. The fluorescent channel on the right corresponds to the magnified inset boxed on the left (white). Note that the movie corresponds to the still images shown in Figure 1A-C and the original movie is re-played once again as marked version, dynamically displaying the line scan data of the VASP signal plus colored arrowheads essentially as shown in the corresponding Figure images. Scale bar corresponds to 10 µm.

**Video S2.** Epifluorescence time-lapse movie of representative B16-F1 WT cell transiently expressing EGFP-VASP and treated with local CD (high dose, 25 µM needle concentration, active treatment indicated with +CD). Cell corresponds to data shown in Figure 1A-C. Scale bar corresponds to 5 µm.

**Video S3.** Epifluorescence time-lapse movie of representative B16-F1 WT cell transiently expressing EGFP-VASP, treated with local CD (low dose, 2 µM needle concentration, active treatment indicated with +CD). Data correspond to the representative example shown in Figure 1A-C. Note that the movie is re-played as marked version highlighting that the reduction of EGFP-VASP signal at the lamellipodial tip upon CD exposure is accompanied by signal movement from tip to rear (yellow arrowheads). Scale bar corresponds to 5 µm.

**Video S4.** Representative time-lapse movies of B16-F1 WT (top row), FMNL2/3 KO (clone #44/3; bottom left) and EVM KO (#23.7.66; bottom right) cells transiently expressing EGFP-VASP (left) or FMNL2-EGFP (right) and transiently treated with local CB (high dose, indicated by yellow ‘+CB’) as indicated and described before. Movies correspond to data shown in Figure 2A-C. Scale bar: 5 µm.

**Video S5.** FRAP movies of B16-F1 cells expressing EGFP-VASP without (left panel) or with (right panel) local application of CB (+CB, high dose), as indicated. Cyan rectangles mark respective bleaching areas. Movies correspond to still images shown in Figure 2D. Scale bar: 5 µm.

**Video S6.** Two consecutive fluorescence time-lapse movies of B16-F1 WT cells transiently expressing EGFP-Fascin or EGFP-MyoX, as indicated, and treated with local CB (high dose, active treatment indicated with ‘+CB’ in each case). Data correspond to Figure 3. Scale bars: 10 µm.

**Video S7.** Two consecutive fluorescence time-lapse movies of B16-F1 WT cells transiently expressing EGFP-β-actin and subjected to CB treatment (high dose, active treatment indicated with ‘+CB’), followed by a combination of two FRAP movies with (right) and without (left) local CB application as control. Details of FRAP movies as described for Video S5. Movies correspond to data displayed in Figure 4A as well as still images shown in Figure 4B. Scale bars: 5 µm.

**Video S8.** Epifluorescence time-lapse movie of a B16-F1 WT cell transiently expressing EGFP-Abi1 and treated with local CB (high dose, indicated by yellow ‘+CB’). Data correspond to the kymograph (left panel) in Figure 4C. Scale bars: 5 µm.

**Video S9.** Epifluorescence time-lapse movies of B16-F1 WT, FMNL2/3 KO (clone #44/3) and EVM KO (clone #23.7.66) cells transiently expressing EGFP-p16B (ArpC5L) and treated with local CB (high dose, indicated by yellow ‘+CB’). Data correspond to kymographs shown in Figure 4D. Scale bars are 5 µm.

**Video S10.** Representative epifluorescence movies of B16-F1 WT, FMNL2/3 KO (#44/3) and EVM KO (clone #23.7.66) cells transiently expressing EGFP-CPβ2 and treated with local CB (high dose, indicated by yellow +CB). Data correspond to kymographs shown in Figure 4E. Scale bars are 5 µm.

**Video S11.** Consecutive epifluorescence movies of first, a B16-F1 CP KO (clone #10) cell expressing EGFP-VASP followed by a B16-F1 CP KO cell co-expressing mCherry-VASP and EGFP-CPβ2 (rescue), and each treated with local CB (high dose, indicated by yellow +CB). The first movie corresponds to Figure 5A, B and C (left panel), and the second to Figure 5C (right panel). Note the rescue of VASP accumulation upon CB application in case of co-expression of CPβ2. Scale bar: 5 µm.

**Video S12.** Phase contrast and epifluorescence video microscopy of a B16-F1 CP KO (clone #10) cell expressing EGFP-VASP and treated with high dose local CD (indicated by yellow +CD). Scale bar: 10 µm.

**Video S13.** Representative time-lapse movies of B16-F1 CP KO (clone #10) cells transiently expressing EGFP-Abi1 (left) and EGFP-β-actin (right) and treated with local CB (high dose, indicated by yellow ‘+CB’) as indicated and described before. Movies correspond to the kymographs (middle and right panels) shown in Figure 6B. Scale bar: 5 µm.

